# Discovery of Small-Molecule Antagonists of the PWWP Domain of NSD2

**DOI:** 10.1101/2020.11.25.398586

**Authors:** Renato Ferreira de Freitas, Yanli Liu, Magdalena M. Szewczyk, Naimee Mehta, Fengling Li, David McLeod, Carlos Zepeda-Velázquez, David Dilworth, Ronan P. Hanley, Elisa Gibson, Peter J. Brown, Rima Al-Awar, Lindsey Ingerman James, Cheryl H. Arrowsmith, Dalia Barsyte-Lovejoy, Jinrong Min, Masoud Vedadi, Matthieu Schapira, Abdellah Allali-Hassani

**Affiliations:** Structural Genomics Consortium, University of Toronto, Toronto, Ontario M5G 1L7, Canada; Center for Integrative Chemical Biology and Drug Discovery, Division of Chemical Biology and Medicinal Chemistry, UNC Eshelman School of Pharmacy, University of North Carolina at Chapel Hill, Chapel Hill, North Carolina 27599, United States; Drug Discovery Program, Ontario Institute for Cancer Research, Toronto, ON M5G 0A3, Canada; Princess Margaret Cancer Centre and Department of Medical Biophysics, University of Toronto, Toronto, ON, M5G 2M9, Canada; Department of Physiology, University of Toronto, Toronto, ON M5S 1A8, Canada; Department of Pharmacology & Toxicology, University of Toronto, Toronto, Ontario M5S 1A8, Canada; Centro de Ciências Naturais e Humanas, Universidade Federal do ABC, Rua Arcturus 3, São Bernardo do Campo, SP 09606-070, Brazil; College of Pharmaceutical Sciences, Soochow University, Suzhou, Jiangsu 215123, China; Incyte, 1801 Augustine Cut-Off, Wilmington, DE 19803, USA

## Abstract

Increased activity of the lysine methyltransferase NSD2 driven by translocation and activating mutations is associated with multiple myeloma and acute lymphoblastic leukemia, but no NSD2-targeting chemical probe has been reported to date. Here, we present the first antagonists that block the protein-protein interaction between the N-terminal PWWP domain of NSD2 and H3K36me2. Using virtual screening and experimental validation, we identified the small-molecule antagonist **3f**, which binds to the NSD2-PWWP1 domain with a K_d_ of 3.4 μM and abrogates histone H3K36me2 binding in cells. This study establishes an alternative approach to targeting NSD2 and provides a small-molecule antagonist that can be further optimized into a chemical probe to better understand the cellular function of this protein.

## INTRODUCTION

NSD2 (nuclear receptor-binding SET domain-containing 2, also known as WHSC1 and MMSET) is a protein lysine methyltransferase that belongs to the NSD family, which also includes NSD1 and NSD3, and that predominantly mono- and dimethylates lysine 36 of histone 3 (H3K36)^1^. NSD2 is an oncoprotein that is aberrantly expressed, amplified or somatically mutated in multiple types of cancer.^2^ Notably, the t(4;14) NSD2 translocation in multiple myeloma (MM) and the hyperactivating NSD2 mutation E1099K in a subset of pediatric acute lymphoblastic leukemia (ALL) result in altered chromatin methylation that drives oncogenesis.^3–5^

While NSD2 is an attractive therapeutic target, efforts to target the catalytic SET domain with small molecule inhibitors have so far met with little success^6–9^ and only recently was the first selective inhibitor of an NSD family protein reported, a compound that binds covalently to the catalytic site of NSD1^10^. Aside from the catalytic domain, NSD2 has multiple protein-protein interaction (PPI) domains that may be clinically relevant, including PHD (plant homeodomain) and PWWP (proline-tryptophan-tryptophan-proline) domains.^11,12^ The N-terminal PWWP domain of NSD2 (NSD2-PWWP1) binds H3K36me2, presumably through a conserved aromatic cage; the F266A mutation at the aromatic cage destabilizes chromatin occupancy of full-length NSD2 and inhibits cancer cell proliferation, but without significantly affecting H3K36 dimethylation.^12^ Small molecules selectively targeting the aromatic cage of NSD2-PWWP1 would be valuable chemical tools to probe the therapeutic relevance of this domain in NSD2-driven tumors, or to design NSD2-targeting PROTACs.

In this study we report the first chemical antagonists targeting the NSD2-PWWP1:H3K36me2 interaction in biophysical and cellular assays. X-ray crystallography established the structural determinants for inhibitor binding and provided crucial information for future ligand optimization. To our knowledge, the present study is the first successful example of using a virtual screening approach to discover antagonists that directly and specifically target a PWWP domain.

## RESULTS AND DISCUSSION

### Structure-based discovery of NSD2-PWWP1 ligands

This project was initiated by the structure-based virtual screening of a library ∼2 million commercial compounds against the PWWP domain of ZMYND11.^13^ A total of 39 compounds were purchased and experimentally screened using differential static light scattering (DSLS – Table S1). Since no hit was confirmed for ZMYND11, we took a target class approach and screened the purchased compound set against other members of the PWWP family. Compound **1** (Figure 1) displayed a stabilization effect on the N-terminal PWWP domain of NSD2, increasing its melting temperature by 4 °C at 400 µM. The interaction of **1** with NSD2-PWWP1 was further confirmed by surface plasmon resonance (SPR), which yielded a dissociation constant (K_d_) of 41 ± 8 μM (Figure S1a). Given the poor solubility of **1**, a reverse isothermal titration calorimetry (ITC) titration was carried out by titrating a concentrated NSD2-PWWP1 solution (1 mM) into the sample cell containing a solution of compound **1** at 40 μM, which produced a K_d_ of 8.9 μM (Figure S1b). Compound **1** did not bind six other PWWP-containing proteins when tested at 400 μM using DSLS (Table S2), indicating that this compound is selective for NSD2-PWWP1. Compound **1** contains two chiral centers but was found to be a single diastereoisomer where both enantiomers bound the target equipotently (Figures S2-4).

**Figure 1.**
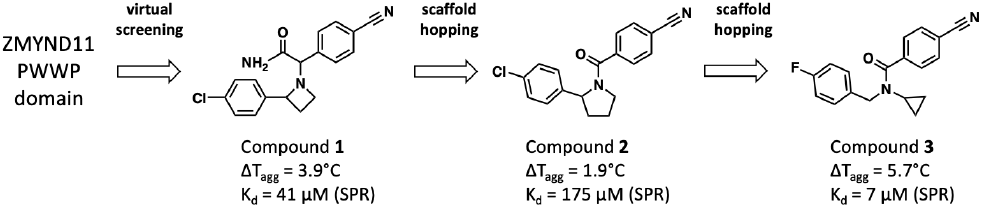
Chemical evolution of NSD2-PWWP1 antagonists. Receptor-based virtual screening followed by target and scaffold hopping led to development of **3**.

### Ligand-guided optimization leads to improved compounds

The encouraging potency and selectivity of **1** prompted us to further explore this scaffold. However, no analogs were commercially available, and only a limited number of building blocks could be purchased for analog synthesis. We therefore sought to identify a new series of antagonists that would be more tractable for follow-up medicinal chemistry efforts. Since no structure of the NSD2-PWWP1 domain had been reported, we used ligand-based scaffold hopping approaches.

First, a low energy 3D conformation of compound **1** was generated with ICM^14^ and used as a reference to screen a virtual library of ∼8 million commercially available compounds with the shape-matching tool ROCS (OpenEye).^15^ Second, we used FTrees (BioSolveIT)^16^ to perform a 2D similarity search against the same library. This approach uses a fuzzy similarity searching, ignoring the three-dimensional structure and chirality of the reference. This feature was helpful in this case, as at the time, we knew neither the bioactive conformation of **1**, nor the correct stereochemistry of the two chiral centers. In the third approach, we performed a substructure search using Filter (OpenEye)^17^. Finally, based on the structure of **1**, a series of scaffold-hopping candidates was manually designed and the closest commercial compounds selected. The atomic property fields (APF)^18^ alignment method implemented in ICM was used to flexibly align the top hits of each approach to the lowest energy conformation of **1** used in the ROCS screen. Visual analysis and clustering with LibMCS (ChemAxon)^19^ led to the selection of 24 compounds (Figure 1, Table S1).

Of the 24 compounds tested, only one hit (compound **2**) showed a modest stabilization (Δ_Tagg_ = 1.9 °C) of NSD2-PWWP1 (Figure 1). The binding of **2** was further confirmed by SPR with a K_d_ of 175 μM. Although **2** was weaker than the initial hit, this new scaffold allowed us to perform a SAR by catalog, as several analogs were commercially available. From a substructure search, 34 analogs of **2** were purchased and tested (Table S1). The initial SAR showed that removing the chlorine group improved potency (**2a**), while replacing it with a methoxy group had no significant effect (**2b**, Table 1). The chlorophenyl group could be replaced with other heterocycles without a significant effect on potency (**2c**-**2e**) but removing the nitrile or replacing it with groups lacking a hydrogen bond acceptor abolished activity (Table 1: **2h vs 2a, 2i vs 2**), suggesting that the nitrile group was forming a specific interaction with the protein.

**Table 1.**
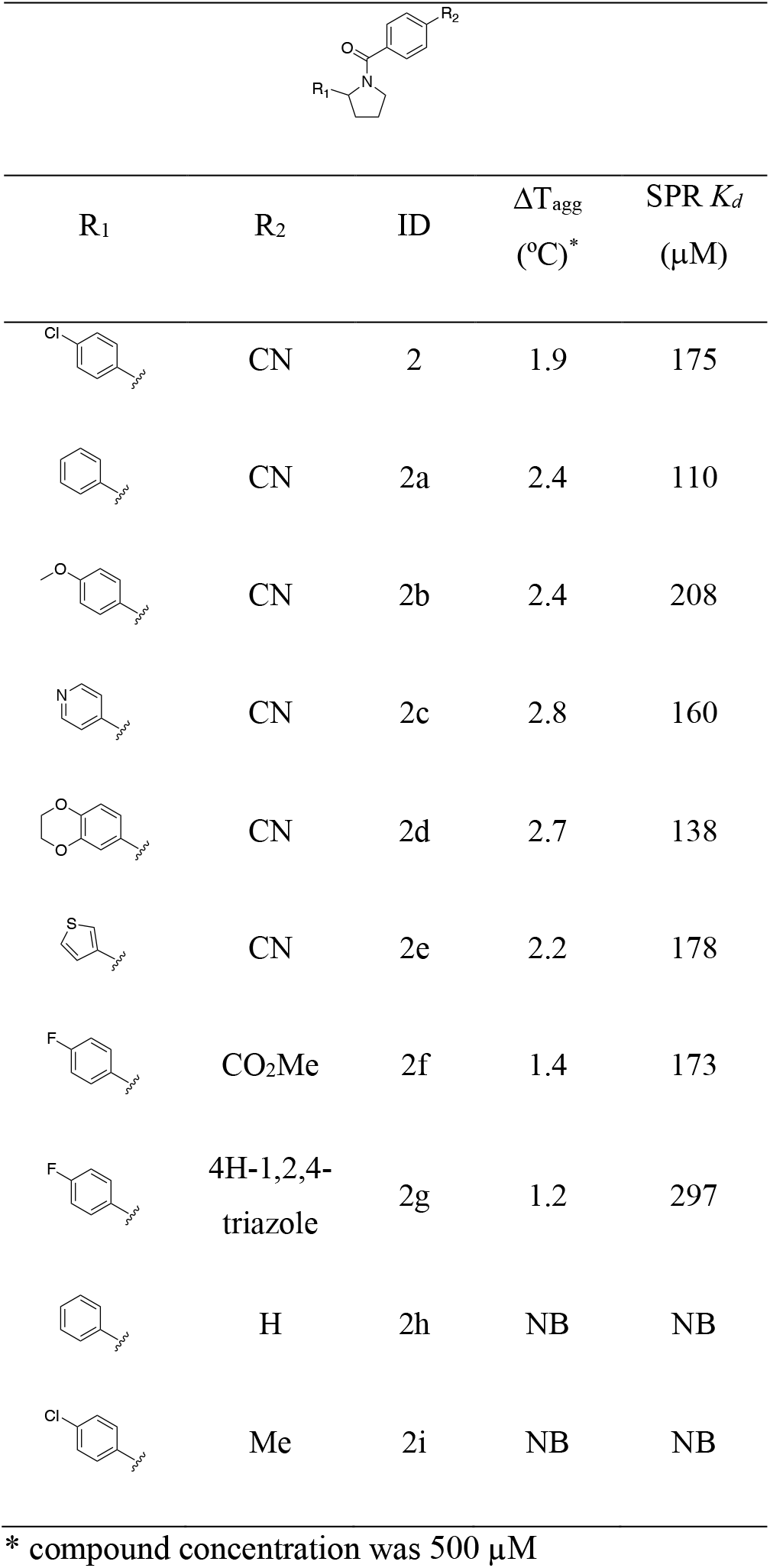
Structure-activity relationships (SAR) of the analogues of compound **2** and close analogues.

|  |  |  |  |  |
| --- | --- | --- | --- | --- |
| R<sub>1</sub> | R<sub>2</sub> | ID | $\Delta T_{agg}$<br>(°C)* | SPR $K_d$<br>(μM) |
|  | CN | 2 | 1.9 | 175 |
|  | CN | 2a | 2.4 | 110 |
|  | CN | 2b | 2.4 | 208 |
|  | CN | 2c | 2.8 | 160 |
|  | CN | 2d | 2.7 | 138 |
|  | CN | 2e | 2.2 | 178 |
|  | CO<sub>2</sub>Me | 2f | 1.4 | 173 |
|  | 4H-1,2,4-triazole | 2g | 1.2 | 297 |
|  | H | 2h | NB | NB |
|  | Me | 2i | NB | NB |

\* compound concentration was 500 μM

On the basis of the preliminary SAR provided by **2** and its close derivatives, we set out to identify novel chemical templates that would preserve the two phenyl groups and the nitrile of **2** using FTrees. The seven compounds selected from this exercise (Table S1) included the break-through antagonist **3**, which binds NSD2-PWWP1 with a K_d_ of 7 μM by SPR and shows strong protein stabilization (Δ_Tagg_ = 5.7 °C) by DSLS (Figure 1). Compound **3** is 6 and 25-times more potent than **1** and **2** respectively, is chemically more accessible than the previous scaffolds as it has no chiral center and can easily be derivatized.

To further understand the molecular basis for the increased potency of **3**, we tested 40 analogs that were either purchased or synthesized (Table 2, Table S1). The initial SAR demonstrated that the cyclopropyl group is an essential structural element, since compounds where it is absent (**3b**) or is replaced with smaller (**3a**) or larger (**3c** – **3e**) groups are much weaker or completely inactive (Table 2). Moreover, replacing the fluorophenyl group with thiophene (**3f**) or tetrahydrofuran (**3g**) resulted in similar potency (Table 2). When tested against 10 PWWP domains by DSLS, compound **3f** only bound NSD2-PWWP1 and was found to be about six times more potent than the initial compound **1** (SPR K_d_ = 7 ± 3 µM) (Figure 2). **3f** did not bind to NSD2-PWWP1 mutants where aromatic cage residues Y233 or F266 were mutated to alanine, indicating binding at the methyl-lysine binding site (Figure S5). In agreement with a recent observation that high DMSO concentrations can limit the potency of PWWP ligands^20^, the K_d_ value of **3f** was measured as 3.4 ± 0.4 µM in an SPR experiment with 0.5% DMSO (Figure S6).

**Table 2.**
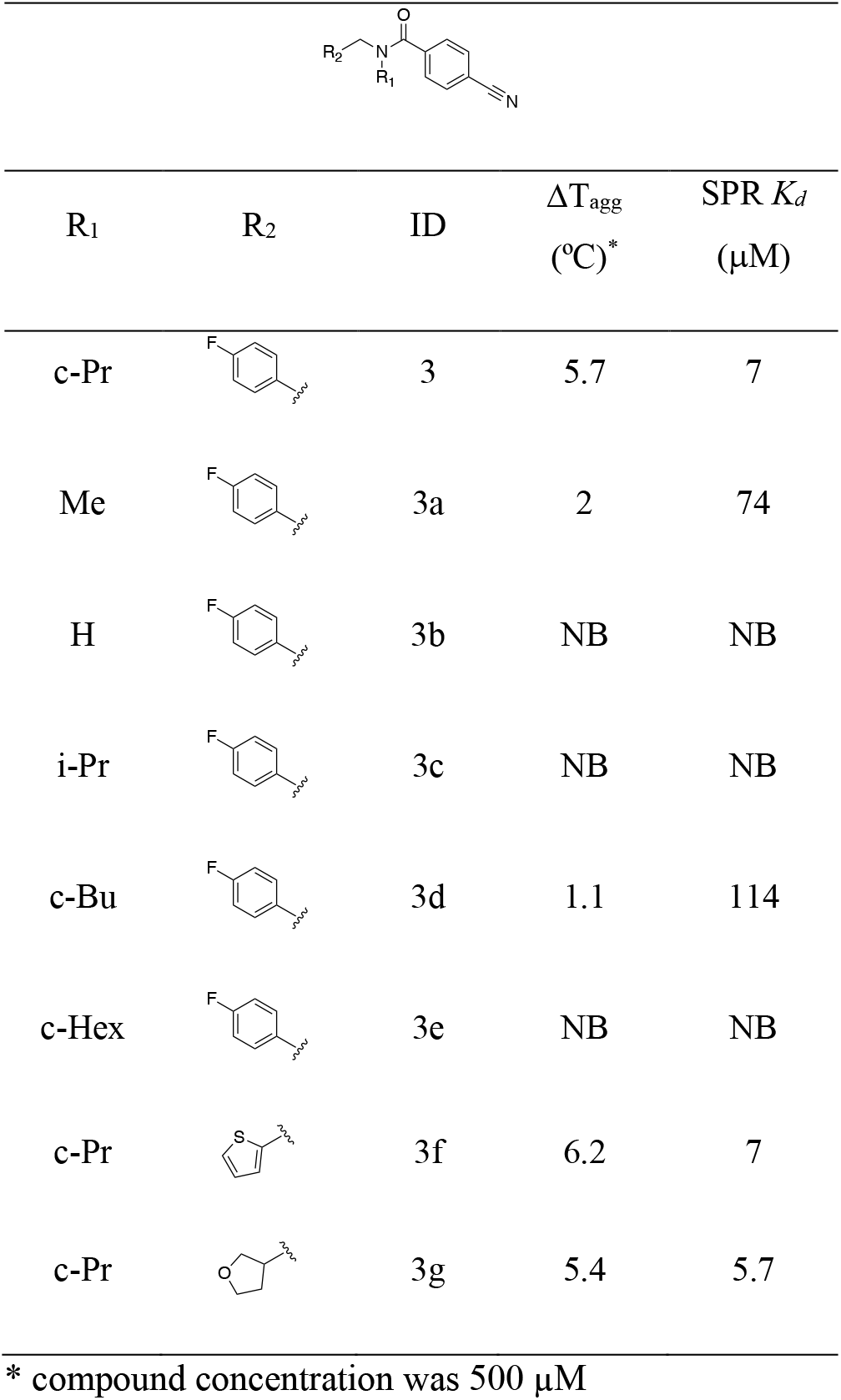
SAR of the analogues of compound **3**

|  |  |  |  |  |
| --- | --- | --- | --- | --- |
| R<sub>1</sub> | R<sub>2</sub> | ID | $\Delta T_{agg}$<br>(°C)* | SPR $K_d$<br>(μM) |
| c-Pr |  | 3 | 5.7 | 7 |
| Me |  | 3a | 2 | 74 |
| H |  | 3b | NB | NB |
| i-Pr |  | 3c | NB | NB |
| c-Bu |  | 3d | 1.1 | 114 |
| c-Hex |  | 3e | NB | NB |
| c-Pr |  | 3f | 6.2 | 7 |
| c-Pr |  | 3g | 5.4 | 5.7 |

\* compound concentration was 500 μM

**Figure 2.**
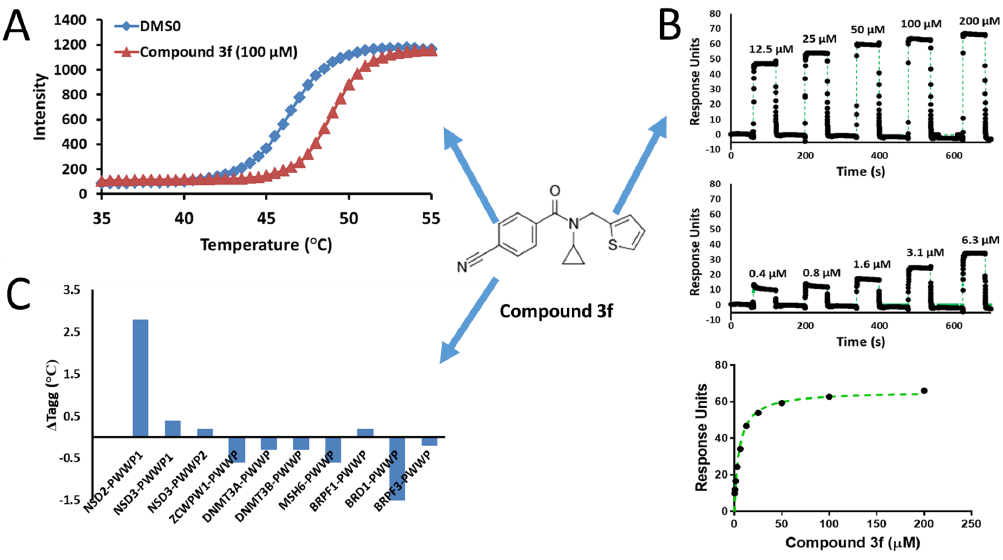
Compound **3f** selectively binds NSD2 PWWP1. (A) treatment with **3f** increases the melting temperature of NSD2-PWWP1 in a DSLS assay. (B) **3f** binds NSD2-PWWP1 in an SPR assay. (C) **3f** only binds to NSD2-PWWP1 in a panel of 10 PWWP domains at 100 µM. The compound structure is shown.

### Crystal structure confirms binding at the PWWP aromatic cage

To support follow-up chemistry focused on this chemical series, we solved the crystal structure of NSD2-PWWP1 in complex with compound **3f** at a resolution of 2.4 Å (Figure 3, Table S3). Examination of the complex structure revealed the molecular basis for the high affinity of the antagonist and confirmed that it was binding in Kme reader aromatic cage. The cyclopropyl ring is bound in a narrow and deep lipophilic pocket formed by the aromatic cage residues (Y233, W236, and F266) and V230, while a tightly coordinated water molecule is found at the bottom of the pocket. This rationalizes previous SAR showing that compounds lacking the cyclopropyl ring are inactive, as they do not fill the hydrophobic pocket. Conversely compounds featuring larger rings such as cyclohexane (**3e**, Table 2), or even an iso-propyl group (**3c**, Table 2) probably clash with the amino acids of the aromatic cage (Figure S7). The cyanophenyl group is enclosed in a pocket formed by W236, G268, D269, A270, E272, L318, and Q321. The nitrile group is completely shielded from the solvent and the nitrogen accepts a hydrogen bond from the amide nitrogen of A270. This supports our previous SAR showing that compounds without the nitrile group are weak or inactive. In addition, the carbonyl oxygen of **3f** is forming a hydrogen bond with the side chain of Y233 (Figure 3). Finally, the thiophene ring is engaged in hydrophobic interactions with V230 and A274, and partially exposed to solvent, which is in agreement with the fact that chemical modifications are tolerated at this end of the molecule.

**Figure 3.**
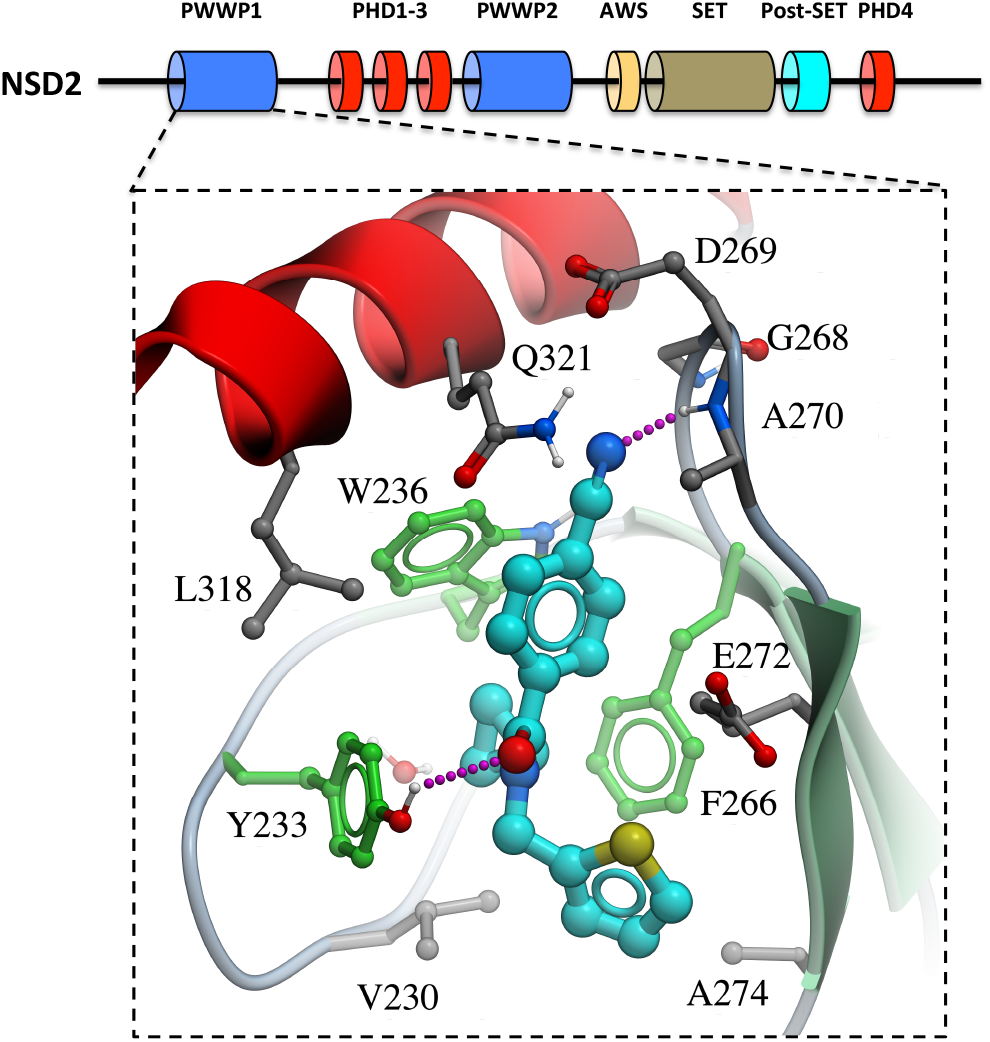
Crystal structure of the NSD2-PWWP1 domain in complex with **3f**. Top: Domain architecture of NSD2. Bottom: Binding pocket residues (grey) and the antagonist (cyan) are displayed as sticks and hydrogen bonds as magenta dashed lines. Aromatic cage residues are colored in green.

Compared to the recently published structure of NSD2-PWWP1 in complex with DNA (PBD ID: 5VC8), the loop residues G268, D269, A270, and P271 connecting the β3 and β4 strands undergo significant conformational changes when the ligand binds (Figure S8). Also, **3f** induces a conformational change of Y233 and E272, opening-up the aromatic cage which was occluded in the apo structure.

### Compound 3f engages NSD2-PWWP1 in cells

Cellular target engagement, and displacement of NSD2-PWWP1 from histone H3.3 was tested using a Nano-BRET assay measuring displacement of NanoLuc-tagged NSD2-PWWP1 domain from Halo-tagged histone H3.3. Encouragingly, we found that compound **3f** decreased interaction of NSD2-PWWP1 but not NSD3-PWWP1 with histone 3 in cells in a dose dependent manner with an IC50 of 17.3 µM (Figure 4).

**Figure 4.**
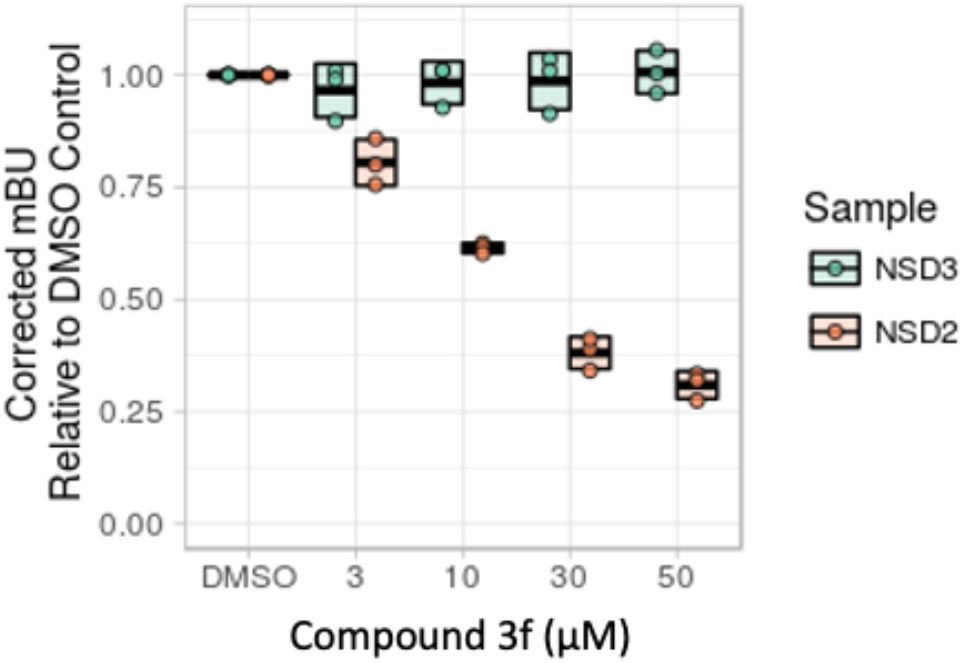
Compound **3f** disengages NSD2 PWWP1 from histone H3 in cells. Mean corrected NanoBRET ratios (mBU) are shown for the interaction between NSD2 (orange) and NSD3 (green) with increasing doses of compound **3f**.

The first potent antagonist for a PWWP domain was recently reported against NSD3-PWWP1^**20**^. Our results further support the idea that PWWP domains represent a chemically tractable target class. PWWP domains are often found in multi-modular proteins involved in transcriptional regulation or DNA repair^**21**^, and pharmacological targeting of PWWP domains could provide an avenue to de-regulate the function of these proteins. PWWP ligands could also serve as chemical handles towards the development of PROTACs that catalyze the proteasomal degradation of PWWP-containing proteins.

## CONCLUSION

In summary, using a combination of receptor-based virtual screening and ligand-based scaffold hopping, we identified chemically tractable small molecule antagonists targeting the NSD2-PWWP1 domain with low micromolar affinity. A crystal structure showed that compound **3f** occupies the Kme binding aromatic cage of NSD2-PWWP1 and **3f** inhibits histone H3K36me2 binding in cells. This study establishes an alternative approach to targeting NSD2 and reveals a class of compounds that we hope to further optimize into a high-quality chemical probe.

## EXPERIMENTAL SECTION

### Computational Methods

#### Docking

The X-ray structure of the PWWP domain of ZMYND11 (PDB ID: 4N4I)^22^ was prepared with PrepWizard (Schrodinger, New York) using the standard protocol, including the addition of hydrogens, the assignment of bond order, assessment of the correct protonation states, and a restrained minimization using the OPLS-AA 2005 force field. Receptor grids were calculated at the centroid of the trimethyllysine (M3L) with the option to dock ligands of similar size.

A lead-like library of ∼2 million compounds was prepared with LigPrep (Schrodinger, New York). The resulting library was then docked using the virtual screening workflow (VSW), using Glide HTVS in the first stage followed by Glide SP in the second stage (Schrodinger, New York). After removing the compounds with a docking score worse than - 7 kcal/mol and that do not contain a tertiary amine, the remaining 18K compounds were docked with Glide XP. Next, compounds with a docking score better than −8 kcal/mol or with a score better than −6 kcal/mol and two hydrogen bonds with nearby residues (R317, F311, H313, N316, W319, H314, S340) were selected for visual inspection. This step selected 431 compounds, which were rescored with the GBSA method (AMBER 12, UCSF). The compounds were clustered and the ones with GBSA > −15 kcal/mol were removed. Finally, after a visual inspection, 40 compounds were selected to be purchased.

#### ROCS search

A library of ∼8 million compounds was processed with the Omega software (OpenEye Scientific Software, Santa Fe) to generate 200 lowest energy conformations for each compound. Then, ROCS (OpenEye Scientific Software, Santa Fe) was used to perform a 3D shape-based search against the resulting library using a low energy 3D conformation of **1** generated by ICM (Molsoft, San Diego) as the query. The compounds were ranked by the TanimotoCombo score, which is a combination of the shape and color similarities, and the top 7500 were saved for analysis.

#### FTrees search

The 2D structure of **1** was used as the reference to search the virtual library described before using the Ftrees (Feature Trees) software (BioSolveIT GmbH)). Feature trees are descriptors that represent the molecule as a reduced graph. The graph encodes functional groups and rings to single nodes. The similarity between two molecules is calculated by comparing their associated feature trees by superposing (matching) similar subtrees onto each other. The top 1980 compounds with the highest similarity with compound 1 were selected for further analysis.

#### Substructure search

The software Filter (OpenEye Scientific Software, Sant Fe) was used to perform a substructure search against the virtual library using a substructure of **1** as the query. The substructure used in the search encoded in SMARTS was NC(=[OX1])C([ar5,ar6])[N;!$(N=C);!$(N=N);!$(N-[!#6;!#1]);!$(N-C=[O,N,S])][CX4][ar5,ar6]. This searched resulted in 1041 compounds.

### Isothermal Titration Calorimetry (ITC)

Purified NSD2-PWWP1 was dialyzed in isothermal titration calorimetry (ITC) buffer (20 mM Tris, pH 7.5, and 100 mM NaCl). Protein at 1 mM was injected into the sample cell containing about 300 μL of 40 μM **1** with 1 % DMSO. ITC titrations were performed on a Nano ITC from TA Instruments (New Castle, DE) at 25 °C by using 4 μL injections with a total of 12 injections. The concentration of DMSO was adjusted to 1% for NSD2-PWWP1 solution. Data were fitted with a one-binding-site model using Nano Analyze Software.

### Differential static light scattering (DSLS)

Experiments were carried out by determining the effect of compounds on the thermal stability by differential static light scattering (DSLS) using StarGazer (from Harbinger). All PWWP domains for selectivity experiments were used at final concentration of 0.2 mg/mL in a buffer consisting of 0.1 M HEPES (pH 7.5) and 150 mM NaCl. As described previously^23^, this method assesses ligand binding through ligand induced stability where protein aggregation/denaturation is measured while heating the sample from 25 to 85 °C at 1 °C/min in a 50 μL volume (covered with 50 μL of mineral oil to prevent evaporation) in a clear-bottom 384-well plate (from Nunc). Aggregation was monitored by the increase of scattered light using CCD camera detection. Pixel intensities were integrated using image analysis software, plotted against temperature and data was then fitted to a Boltzmann sigmoid function to obtain the aggregation temperature (T_agg_) from the midpoint of the transition.

### Surface Plasmon Resonance (SPR)

SPR studies were performed using a Biacore T200 (GE Health Sciences Inc.) at 20 °C. Biotinylated NSD2-PWWP1 was captured onto one flow cell of a streptavidin-conjugated SA chip at approximately 6,000 response units (RU) (according to manufacturer’s protocol) while another flow cell was left empty for reference subtraction. Compound **3f** was tested at 200 μM as the highest concentration and dilution factor of 0.5 in in HBS–EP buffer (20 mM HEPES pH 7.4, 150 mM NaCl, 3 mM EDTA, 0.05% Tween-20) was used to yield 10 concentrations. Experiments were performed using the same buffer with 5% DMSO in single cycle kinetic with 60s contact time and a dissociation time of 120s at a flow rate of 75 µL/min. Kinetic curve fittings and K_d_ value calculations were done with a 1:1 binding model using the Biacore T200 Evaluation software (GE Health Sciences Inc.).

### Crystal structure

The NSD2 fragment (aa 211-350) containing the first human PWWP domain of NSD2 was cloned into a pET28a-MHL vector with a His-tag in its N-terminus for affinity purification. The recombinant plasmid was transformed in the E coli BL21 (DE3) and the target protein overexpression was induced by 0.2 mM IPTG at 18°C overnight. The cells were harvested and lysed by sonication. The cell lysis was first purified by Ni-NTA, and the eluted protein was treated by TEV to remove the His-tag by dialysis against dialysis buffer containing 20 mM Tris pH 7.5, 150 mM NaCl, 1 mM DTT. The sample was then purified by gel filtration chromatography using a Superdex 75 column in a buffer containing 20 mM Tris, pH 7.5, 150 mM NaCl, 1 mM DTT. The peak fractions from the gel filtration column were pooled and concentrated to 20 mg/mL for crystallization.The purified protein was mixed with the ligand at a molar ratio of 1:3 and co-crystallized using the sitting drop vapour diffusion method at 18 °C. The crystals were obtained in a buffer containing 20% PEG 3350, 0.2 M KSCN. Crystals were soaked in a cryoprotectant consisting of 100% reservoir solution and 15% glycerol for data collection. X-ray diffraction data for NSD2 + **3f** was collected at 100K at beamline 08ID-1 of Canadian Light Source (CLS). The data set was processed using the XDS^24^ suite. The structures of NSD2 + **3f** was solved by molecular replacement using PHASER^25^ with PDB entry 5VC8 as search template. Graphics program COOT^26^ was used for model building and visualization. Geometry restraints for the compound refinement were prepared with GRADE (Global Phasing Ltd.) developed at Global Phasing Ltd. Restrained refinement and validation using BUSTER (Global Phasing Ltd.), and MOLPROBITY^27^, respectively.

### NanoBRET assay

U2Os cells were plated in 12-well plates (1×10^5^/well) in DMEM supplemented with 10% FBS, penicillin (100 U/mL) and streptomycin (100 µg/mL). 4 hours after plating cells were transfected with 1μg of histone H3.3-HaloTag®Fusion Vector DNA (Promega) + 0.1 μg of NSD2 PWWP1-NanoLuc® Fusion Vector DNA (C-terminal) (Promega) using X-tremeGENE HP DNA Transfection Reagent (Sigma), following manufacturer instructions. Next day cells were trypsinized, spun down and resuspended in DMEM/F12 (no phenol red) supplemented with 4% FBS, penicillin (100 U/mL) and streptomycin (100 µg/mL) at density 1.1×10^5^/mL. Cells were divided into two pools. To the first pool 1 µl/mL DMEM/F12 (no phenol red) of HaloTag® NanoBRET™ 618 Ligand (Promega) was added and to the second pool DMSO. Cells were plated (90 μL/well) in 96-well plates (white, 655083, Greiner Bio One). 10 x concentrated compound and DMSO control was prepared in DMEM/F12 and added to cells (10 μL/well). Next day 25 μL of NanoBRET™ Nano-GloR Substrate (Promega) solution in DMEM/F12 (10 μL/mL) was added to each well. Cells were shaken for 30 s and donor emission at 450 nm (filter: 450 nm/BP 80nm) and acceptor emission at 618 nm (filter: 610nm/LP) was measured within 10 minutes of substrate addition using CLARIOstar microplate reader (Mandel). Mean corrected NanoBRET ratios (mBU) were determined by subtracting mean of 618/460 signal from cells without NanoBRET™ 618 Ligand x 1000 from mean of 618/460 signal from cells with NanoBRET™ 618 Ligand x 1000.

## ASSOCIATED CONTENT

Supporting Information.

Experimental Details, supporting tables and figures, X-ray crystallographic statistics (PDF). The crystallographic coordinates are deposited in the Protein Data Bank (PDB code 6UE6).

## AUTHOR INFORMATION

### Author Contributions

Experimental design: RFF, AAH, MS, MV, NM, LIJ, YL, JM, MMS, DD, DB; data generation: RFF, AAH, NM, YL, FL, DM, MMS, DD, RPH, EG, DML, CZ-V; data analysis: RFF, AAH, MS, MV, NM, LIJ, MMS, DD, DB, PJB, CHA;

### Supervision

MS, MV, JM, LIJ, DB, CZ; manuscript writing: RFF, MS, AAH, DD, MV, LIJ; Funding: CHA, LIJ, RA-A; manuscript review: all

### Notes

The authors declare no competing financial interest.

## ACKNOWLEDGMENT

The SGC is a registered charity (no. 1097737) that receives funds from AbbVie, Bayer Pharma AG, Boehringer Ingelheim, Canada Foundation for Innovation, Eshelman Institute for Innovation, Genome Canada through Ontario Genomics Institute, Innovative Medicines Initiative (EU/EFPIA) [ULTRA-DD grant no. 115766], Janssen, Merck & Co., Novartis Pharma AG, Ontario Ministry of Economic Development and Innovation, Pfizer, São Paulo Research Foundation-FAPESP, Takeda, and the Wellcome Trust. Additional funding from the Collaborative Research and Training Experience grant (Aled Edwards 432008-2013) from the Natural Sciences and Engineering Research Council of Canada. We thank BioSolveIT GmbH for supplying an evaluation copy of the Ftrees. We are grateful to OpenEye Scientific Software, Inc. for providing us with an academic license for their software. MS gratefully acknowledges funding from NSERC (grants RGPIN-2019-04416 and ALLRP 555329-20). This work was further supported by the National Cancer Institute, NIH (grant R01CA242305 to L.I.J.).

---

SYNOPSIS TOC (Word Style “SN_Synopsis_TOC”). If you are submitting your paper to a journal that requires a synopsis graphic and/or synopsis paragraph, see the Instructions for Authors on the journal’s homepage for a description of what needs to be provided and for the size requirements of the artwork.

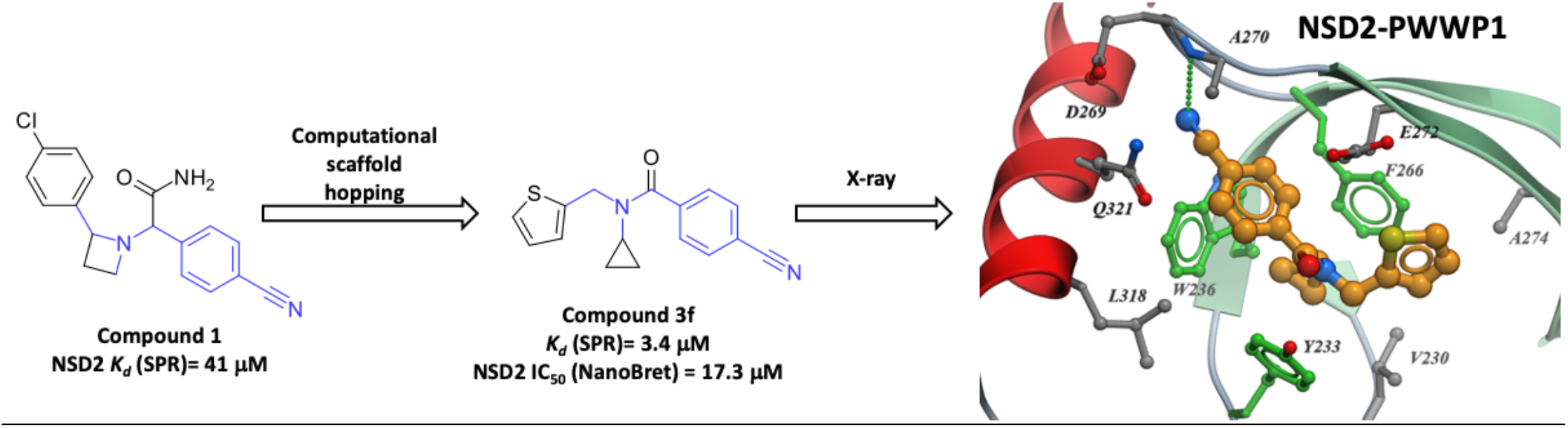

## Notes

### Competing Interest Statement

The authors have declared no competing interest.

