## Supplementary information for "Discovery of Small-Molecule Antagonists of the PWWP Domain of NSD2"

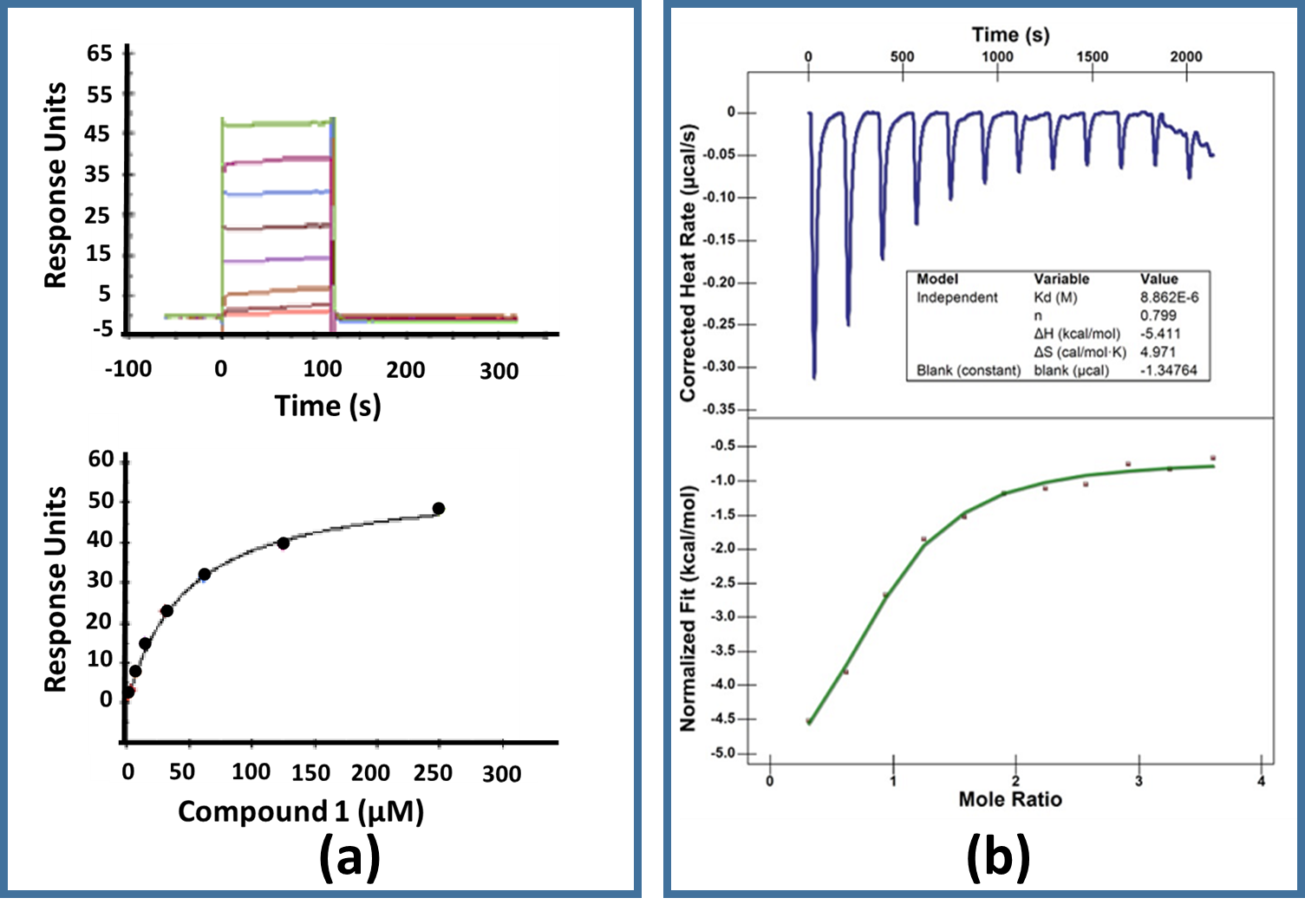

Figure S1. (a) Compound **1** binds NSD2-PWWP1 with a K_d_ of 41 ± 8 µM in SPR experiments conducted in triplicate. SPR-sensorgram (upper pane) for the interaction of **1** with NSD2-PWWP1 domain. Multi-cycle kinetics was used with concentrations from 2 µM to 250 µM (dilution factor of 0.5 was used to yield 8 concentrations). 90 s contact time and 120 s dissociation time at 100 µL/min were used. The steady state values were determined and plotted as a function of the concentration (lower pane). A single binding site model was fitted to the data to calculate K_d_. (b) ITC results showing raw data after integration baseline correction (upper pane) and integrated data and regression (lower pane). n is the number of molecules per binding site, K_d_ is the association constant, ΔH is the change in enthalpy, and ΔS is the change in entropy.

Table S2. Thermal shift in differential static light scattering (DSLS) induced by **1** against seven PWWP domains.

| **Protein** | **ΔT_agg_ at 400 µM (ºC)** |
| --- | --- |
| BRPF1-PWWP | -0.5 |
| DNMT3b-PWWP | -1.2 |
| MSH6-PWWP | -1.4 |
| NSD2-PWWP1 | 3.9 |
| NSD3-PWWP1 | -0.3 |
| ZMYND11-PWWP | -1.2 |
| ZCWPW1-PWWP | -1.4 |

**Characterization of Compound 1**

The original sample was obtained from Enamine (Catalog Number Z1483746373) and assumed to be a racemic mixture. Attempted separation of all 4 diastereoisomers by chiral SFC resulted in only two peaks in the SFC chromatogram. Peaks showed identical NMR spectra but exhibited equal and opposite optical rotations. These must be enantiomers and thus the original sample must have been made by a diasteroselective route (Figure S2).

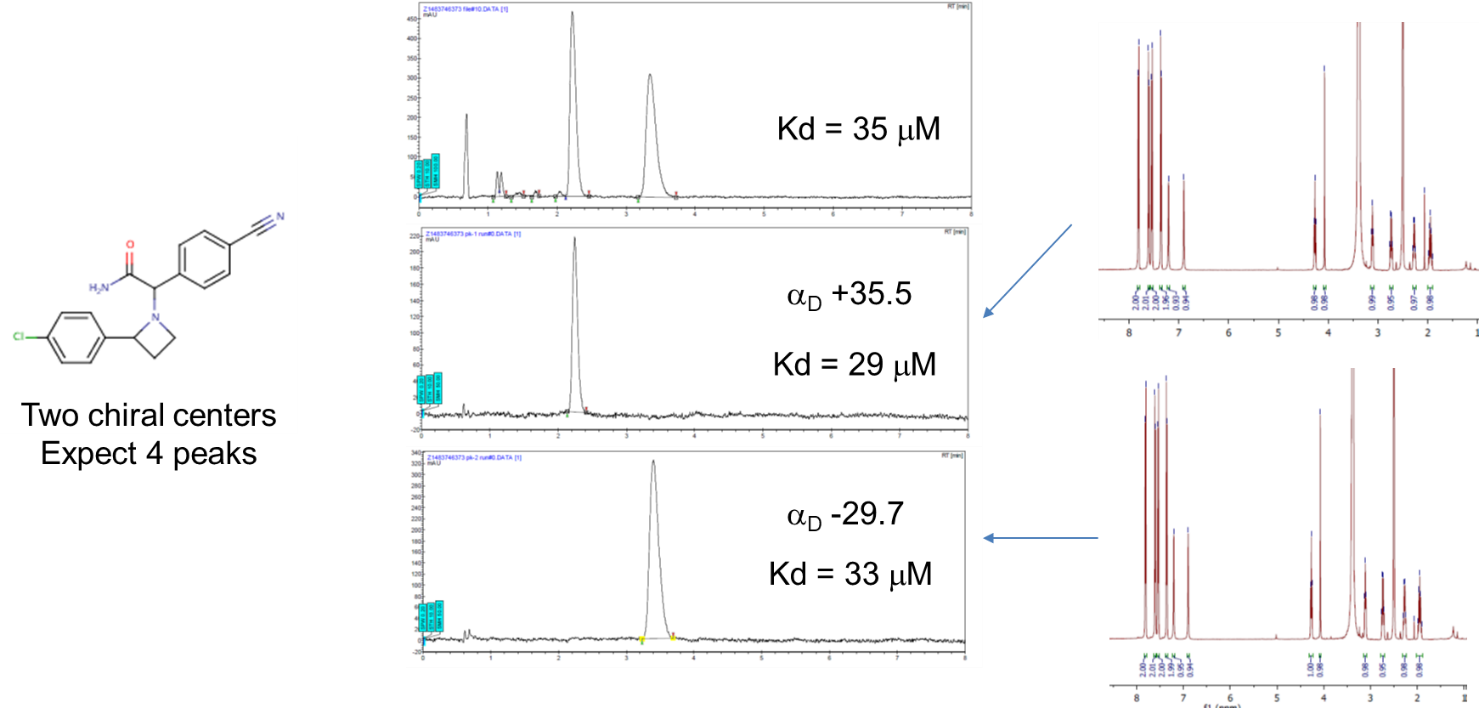

Figure S2. Separation of enantiomers from original sample by chiral SFC.

A search of the literature found a paper describing this diastereoselective synthesis from a substituted boronic acid and cyclic amine (Nanda K. K. and Trotter B. W. **Tet. Lett.** (2005) 2025-28). In an effort to determine the activity of the other diasteroisomers, a non-diastereoselective route was developed involving azetidine displacement of an a-bromoester (Figure S3).

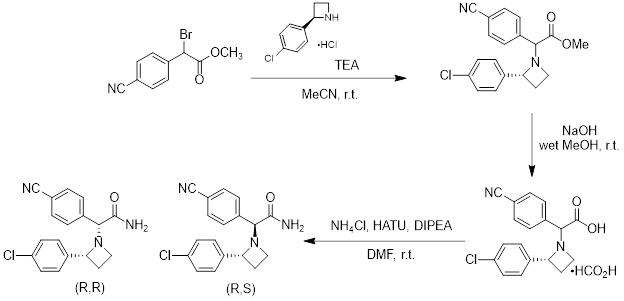

Figure S3. Synthesis from (*R*)-azetidine gave two diastereoisomers. Similar results were obtained starting from (*S*)-azetidine to give the *(S,S*) and (*S,R*) diastereoisomers.

In this case, all four isomers were detected by chiral SFC, however, the diastereoisomers not found in the original material were inactive. The assignment of stereochemistry was based on the diastereoselective product in the original paper which was confirmed by X-ray crystallography (Figure S4).

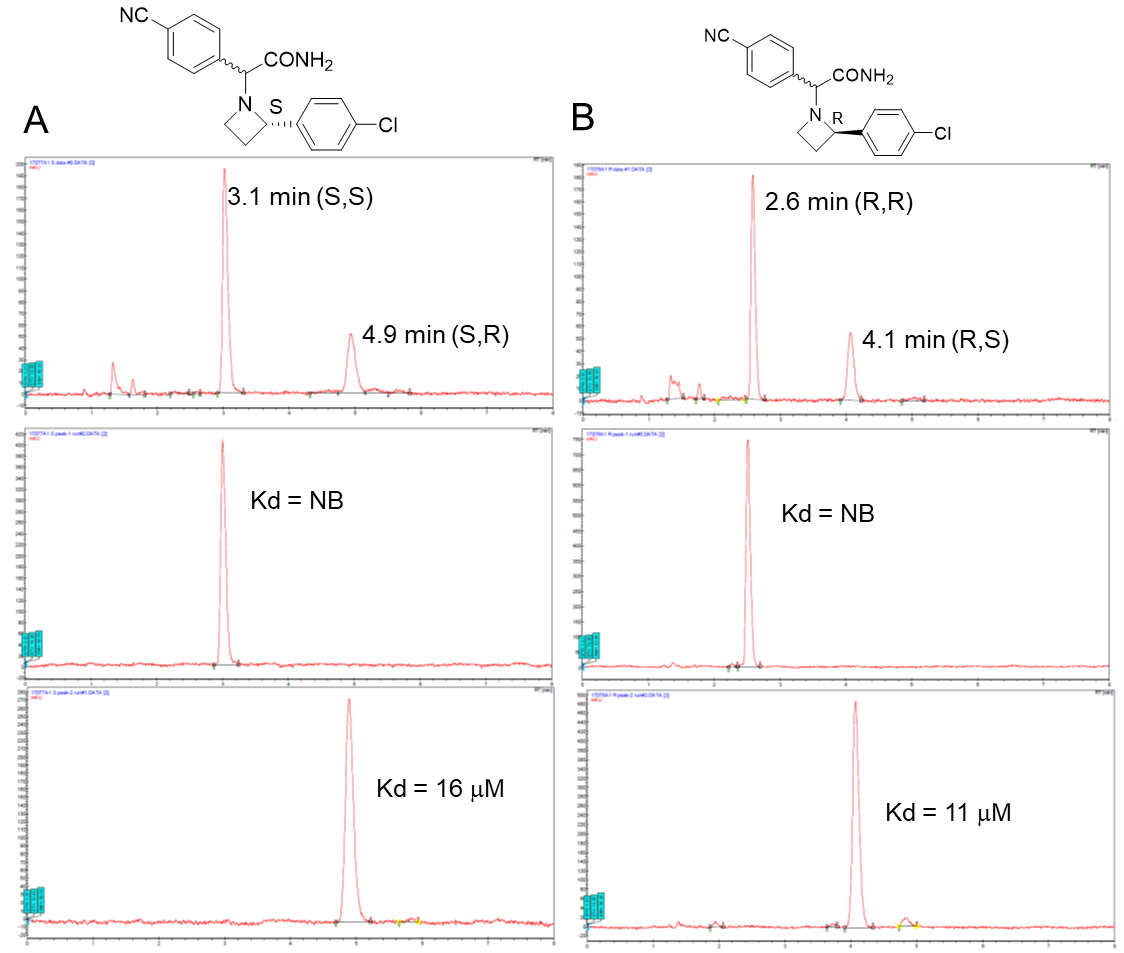

Figure S4. Chiral preparative SFC separation of diastereoisomers. A) Reaction products from (*S*)-azetidine gave peaks at 3.1 and 4.9 mins. B) Products from (*R*)-azetidine gave peaks at 2.6 and 4.1 mins. The original enantiomers correspond to the peaks at 4.1 and 4.9 mins. Lower panels indicate the analytical chiral SFC analysis of collected peaks.

.

**Experimental Procedures and Characterization data**

**Separation of commercial material**

The commercially available material (30 mg; Enamine Z1483746373) was separated by chiral SFC using an IA column (2 x 25 cm) using 30% MeOH/CO_2_ as eluant at a flowrate of 60 mL.min^-1^ and 100 bar pressure. Detection by UV at 220 nm. Injection volume was 1.5 mL of 3 mg.mL^-1^ in MeOH. Analytical chiral SFC (Fig. 1) used IA column (0.46 x 25 cm) with 40% MeOH/CO_2_ as eluant at a flowrate of 3 mL.min^-1^ and 100 bar pressure. Detection by UV at 220 nm.

**2R-(2-(4-chlorophenyl)azetidin-1-yl)-2S-(4-cyanophenyl)acetamide** (13 mg). []_d_^23^ +35.5° (c=0.205; MeOH). ^1^H NMR (500 MHz, DMSO) δ 7.81 (d, *J* = 8.2 Hz, 2H), 7.60 (d, *J* = 8.2 Hz, 2H), 7.54 (d, *J* = 8.4 Hz, 2H), 7.36 (d, *J* = 8.4 Hz, 2H), 7.21 (s, 1H), 6.90 (s, 1H), 4.27 (t, *J* = 8.1 Hz, 1H), 4.07 (s, 1H), 3.11 (t, *J* = 6.5 Hz, 1H), 2.73 (dd, *J* = 16.1, 8.0 Hz, 1H), 2.27 (q, *J* = 8.0 Hz, 1H), 1.95 (p, *J* = 9.0 Hz, 1H).

**2S-(2-(4-chlorophenyl)azetidin-1-yl)-2R-(4-cyanophenyl)acetamide** (12 mg). []_d_^23^ -29.7° (c=0.337; MeOH). ^1^H NMR (500 MHz, DMSO) δ 7.81 (d, *J* = 8.2 Hz, 2H), 7.60 (d, *J* = 8.2 Hz, 2H), 7.54 (d, *J* = 8.3 Hz, 2H), 7.36 (d, *J* = 8.3 Hz, 2H), 7.21 (s, 1H), 6.90 (s, 1H), 4.27 (t, *J* = 8.1 Hz, 1H), 4.07 (s, 1H), 3.11 (t, *J* = 6.5 Hz, 1H), 2.73 (dd, *J* = 16.1, 8.0 Hz, 1H), 2.27 (q, *J* = 8.0 Hz, 1H), 1.95 (p, *J* = 9.0 Hz, 1H).

**Synthesis and separation of all four isomers**

All reagents were purchased from commercial vendors and used without further purification. Volatiles were removed under reduced pressure by rotary evaporation or by using the V-10 solvent evaporator system by Biotage^®^. Very high boiling point (6000 rpm, 0 mbar, 56 ^o^C), mixed volatile (7000 rpm, 30 mbar, 36 ºC) and volatile (6000 rpm, 30 mbar, 36 ^o^C) methods were used to evaporate solvents. The yields given refer to chromatographically purified and spectroscopically pure compounds. Compounds were purified using a Biotage Isolera One system by normal phase chromatography using Biotage^®^ SNAP KP-Sil or Sfär Silica D columns (Part No.: FSKO-1107/FSRD-0445) or by reverse-phase chromatography using Biotage^®^ SNAP KP-C18-HS or Sfär C18 D columns (Part No.: FSLO-1118/FSUD-040). If additional purification was required, compounds were purified by solid phase extraction (SPE) using Biotage Isolute Flash SCX-2 cation exchange cartridges (Part No.: 532-0050-C and 456-0200-D). Products were washed with 2 cartridge volumes of MeOH and eluted with a solution of MeOH and NH_4_OH (9:1 v/v). Preparative chromatography was carried out using a Waters 2767 injector with the collector attached to PDA UV/Vis and SQD mass detectors. An XSelect CSH Prep C18 5µm OBD 19 mm x 100 mm (Part No.: 186005421) or Xselect CSH Prep C18 5µm 10 mm x 100 mm (Part No.: 186005415) column was used for purification. Final compounds were dried using the Labconco^TM^ Benchtop FreeZone^TM^ Freeze-Dry System (4.5 L Model). ^1^H and proton-decoupled ^19^F NMRs were recorded on a Bruker Avance-III 500 MHz spectrometer at ambient temperature. Residual protons of CDCl_3_, DMSO-*d*_6_ and CD_3_OD solvents were used as internal references. Spectral data are reported as follows: chemical shift (δ in ppm), multiplicity (br = broad, s = singlet, d = doublet, dd = doublet of doublets, m = multiplet), coupling constants (*J* in Hz) and proton integration. Compound purity was determined by UV absorbance at 254 nm during tandem liquid chromatography/mass spectrometry (LCMS) using a Waters Acquity separations module. All final compounds had a purity of ≥95% as determined using this method**.** Low resolution mass spectrometry (LRMS) was conducted in positive ion mode using a Waters Acquity SQD mass spectrometer (electrospray ionization source) fitted with a PDA detector. Mobile phase A consisted of 0.1% formic acid in water, while mobile phase B consisted of 0.1% formic acid in acetonitrile. Column 1 (Method 1): Acquity UPLC CSH C18 (2.1 x 50 mm, 130 Å, 1.7 µm. Part No. 186005296). The gradient went from 90% to 5% mobile phase A over 1.8 min, maintained at 5% for 0.5 min, then increased to 90% over 0.2 min for a total run time of 3 min. The column was used with the temperature maintained at 25 °C.

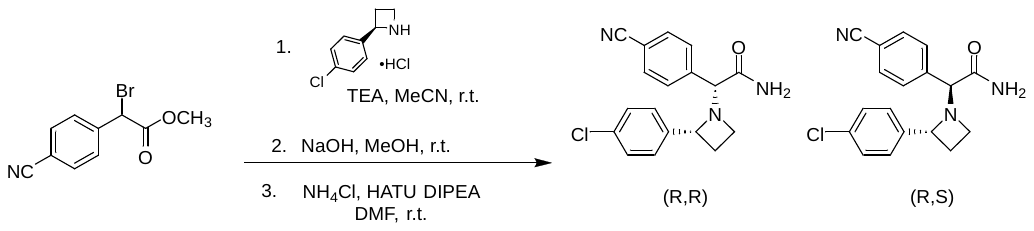

A solution of (*R*)-2-(4-chlorophenyl)azetidine, hydrochloride (67.5 mg, 0.331 mmol) and *N*,*N*-diisopropylethylamine (121 μL, 0.694 mmol) in acetonitrile (0.5 ml) was added dropwise to methyl 2-bromo-2-(4-cyanophenyl)acetate (84 mg, 0.331 mmol) in acetonitrile (1.18 ml) at 0°C. The mixture was allowed to warm to room temperature and stirred for 5 h. All volatiles were then removed under reduced pressure and the residue partitioned between EtOAc and brine. The combined organic layers were then dried over Na_2_SO_4_, concentrated under reduced pressure and column chromatographed (silica gel, hexanes/EtOAc, 10:0 to 7:3 v/v) to afford methyl 2-((R)-2-(4-chlorophenyl)azetidin-1-yl)-2-(4-cyanophenyl)acetate (72 mg, 64 % yield) as a yellow oil. LCMS Method 1, RT = 1.84 and 2.04 min, MS (ESI): *m/z* = 341.3 [M + 1]^+^, Purity (UV^254^) = 99%

t

To the above ester (72 mg, 0.211 mmol) in wet MeOH (10 mL) was added potassium hydroxide (59 mg, 1.06 mmol) and the reaction stirred at room temperature overnight. All volatiles were evaporated and the residue column chromatographed (RP-C18, H_2_O (0.1% v/v FA)/MeCN, 98:2 to 10:90 v/v) to afford 2-2R-((4-chlorophenyl)azetidin-1-yl)-2-(4-cyanophenyl)acetic acid formic acid salt (39 mg, 50 % yield) as a white solid. LCMS Method 1, RT = 1.36 and 1.39 min, MS (ESI): *m/z* = 327.4 [M + 1]^+^, Purity (UV^254^) = 99%

To the above acid salt in DMF (5 mL) were added N,N-diisopropylethylamine (121 μL, 0.694 mmol) and HATU (69.6 mg, 0.183 mmol) at room temperature. To this mixture was added, with vigorous stirring, ammonium chloride (26.1 mg, 0.488 mmol) in one portion and the reaction stirred for 1 h. Water (20 mL) was added to quench the reaction and the aqueous layer was extracted with EtOAc (3 x 10 mL) to give the desired compound as a mixture of diastereomers. LCMS Method 1, RT = 1.52 min, MS (ESI): *m/z* = 326.4 [M + 1]^+^, Purity (UV^254^) = 99%

This synthetic mixture was separated by preparative chiral SFC using an AD-H column (2 x 25 cm) using 35% MeOH/CO_2_ as eluant at a flowrate of 70 mL.min^-1^ and 100 bar pressure. Detection by UV at 220 nm. Injection volume was 1.0 mL of 5 mg.mL^-1^ in MeOH/dichlorometane. Analytical chiral SFC (Fig. 1) used AD-H column (0.46 x 25 cm) with 300% MeOH/CO_2_ as eluant at a flowrate of 3 mL.min^-1^ and 120 bar pressure. Detection by UV at 220 nm.

**2R-(2-(4-chlorophenyl)azetidin-1-yl)-2R-(4-cyanophenyl)acetamide** (29 mg). Chiral SFC retention time 2.6 min. ^1^H NMR (500 MHz, MeOD) δ 7.38 (d, J = 8.3 Hz, 2H), 7.33 (d, J = 8.4 Hz, 2H), 7.12 (d, J = 8.6 Hz, 2H), 7.08 (d, J = 8.6 Hz, 2H), 4.11 (t, J = 8.2 Hz, 1H), 4.07 (s, 1H), 3.73 – 3.69 (m, 1H), 3.13 (dt, J = 9.3, 8.0 Hz, 1H), 2.36 (dtd, J = 10.3, 8.0, 2.3 Hz, 1H), 2.24 – 2.11 (m, 1H).

**2R-(2-(4-chlorophenyl)azetidin-1-yl)-2S-(4-cyanophenyl)acetamide** (13 mg). Chiral SFC retention time 4.1 min. ^1^H NMR (500 MHz, MeOD) δ 7.72 (d, J = 8.3 Hz, 2H), 7.63 (d, J = 8.3 Hz, 2H), 7.54 (d, J = 8.4 Hz, 2H), 7.34 (d, J = 8.4 Hz, 2H), 4.27 (t, J = 8.3 Hz, 1H), 4.11 (s, 1H), 3.27 – 3.20 (m, 1H), 2.89 – 2.80 (m, 1H), 2.32 (dtd, J = 10.4, 7.9, 2.4 Hz, 1H), 2.25 – 2.15 (m, 1H).

Similarly prepared were the diastereoisomers from (*S*)-2-(4-chlorophenyl)azetidine

**2S-(2-(4-chlorophenyl)azetidin-1-yl)-2S-(4-cyanophenyl)acetamide** (29 mg). Chiral SFC retention time 3.1 min. ^1^H NMR (500 MHz, MeOD) δ 7.36 (d, J = 8.3 Hz, 2H), 7.31 (d, J = 8.4 Hz, 2H), 7.10 (d, J = 8.6 Hz, 2H), 7.07 (d, J = 8.6 Hz, 2H), 4.09 (t, J = 8.2 Hz, 1H), 4.05 (s, 1H), 3.72 – 3.66 (m, 1H), 3.11 (dt, J = 9.4, 8.0 Hz, 1H), 2.34 (dtd, J = 10.3, 8.0, 2.3 Hz, 1H), 2.20 – 2.10 (m, 1H).

**2S-(2-(4-chlorophenyl)azetidin-1-yl)-2R-(4-cyanophenyl)acetamide** (7 mg). Chiral SFC retention time 4.9 min. ^1^H NMR (500 MHz, MeOD) δ 7.72 (d, J = 8.4 Hz, 2H), 7.63 (d, J = 8.3 Hz, 2H), 7.54 (d, J = 8.4 Hz, 2H), 7.34 (d, J = 8.4 Hz, 2H), 4.27 (t, J = 8.3 Hz, 1H), 4.11 (s, 1H), 3.28 – 3.20 (m, 1H), 2.89 – 2.80 (m, 1H), 2.32 (dtd, J = 10.4, 8.0, 2.4 Hz, 1H), 2.24 – 2.15 (m, 1H).

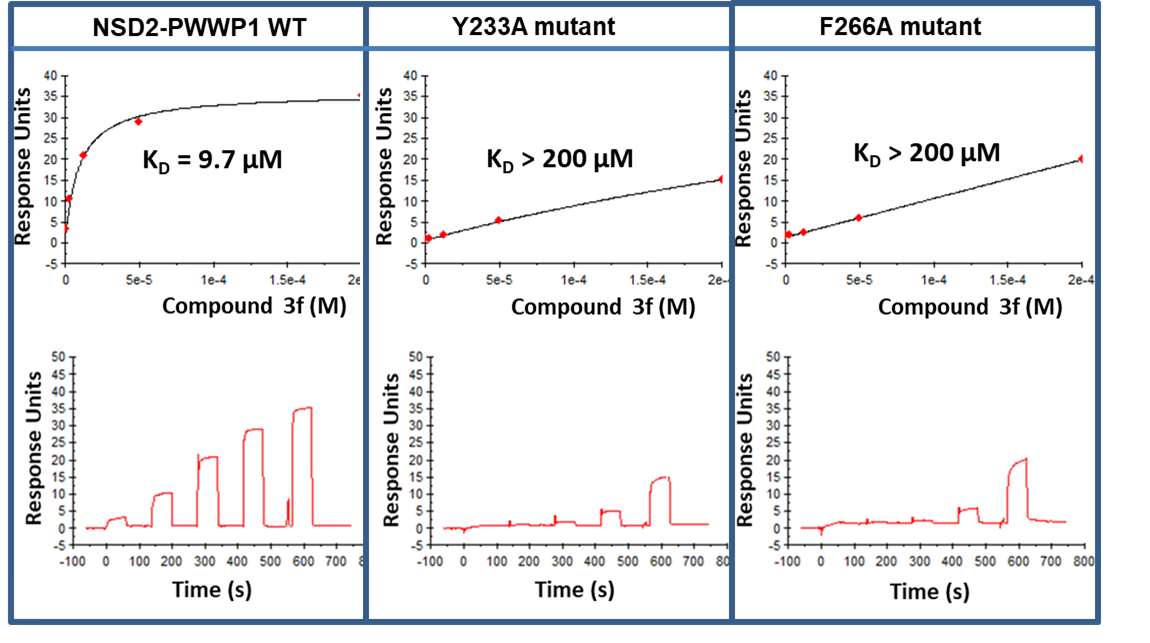

Figure S5: compound **3f** binds wild-type NSD2-PWWP1 but not Y233A or F266A mutants in SPR experiments

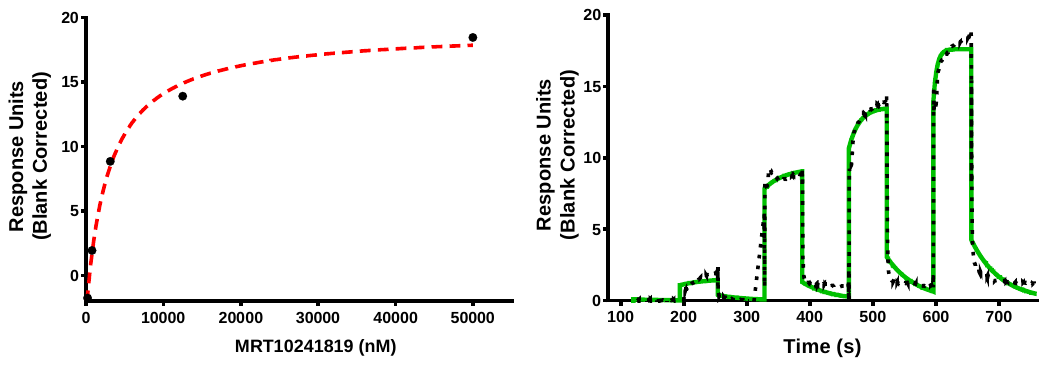

Figure S6. Compound 3f binds NSD2-PWWP1 with a K_d_ of 3.42 ± 0.45 µM in a SPR experiments conducted in triplicate with 0.5% DMSO concentration. Biotinylated NSD2-PWWP1 domain (208- 368) was immobilized on the flow cell of an SA sensor chip in 1x HBS-EP buffer, yielding 5000 RU. The biotinylated RBBP5 (2-538) was immobilized on another flow cell of SA chip, yielding 4300 RU as a negative control. Using the same buffer with 0.5% DMSO and single cycle kinetics with 60 s contact time and a dissociation time of 120s at a flow rate of 75 µL/min. The compound was tested at 50 µM as the highest concentration and dilution factor of 0.25 was used to yield 5 concentrations.

Table S3. Crystallography data and refinement statistics

|  | NSD2 + **3f** |
| --- | --- |
| **PDB Code** | 6UE6 |
| **Data collection** |  |
| Space group | P2_1_2_1_2_1_ |
| Cell dimensions |  |
| *a*, *b*, *c* (Å) | 69.2, 70.3, 228.8 |
| *α*, *β*, *γ* (°) | 90.0,90.0,90.0 |
| Resolution (Å) (highest resolution shell) | 49.33-2.40(2.49-2.40) |
| Measured reflections | 287140 |
| Unique reflections | 44525 |
| *R*_merge_ | 8.4(0.997) |
| *I*/σ*I* | 14.0(1.9) |
| Completeness(%) | 99.7(97.5) |
| Redundancy | 6.4(5.2) |
| **Refinement** |  |
| Resolution (Å) | 45.3-2.40 |
| No. reflections (test set) | 44447(2275) |
| *R*_work/_ *R*_free_ (%) | 23.6/25.2 |
| No. atoms |  |
| Protein | 7136 |
| Compound | 160 |
| B-factors (Å^2^) |  |
| Protein | 61.4 |
| Compound | 47.5 |
| RMSD |  |
| Bond lengths (Å) | 0.010 |
| Bond angles (º) | 0.99 |
| Ramachandran plot % residues |  |
| Favored | 99.5 |
| Additional allowed | 0.5 |
| Generously allowed | 0 |
| Disallowed | 0 |

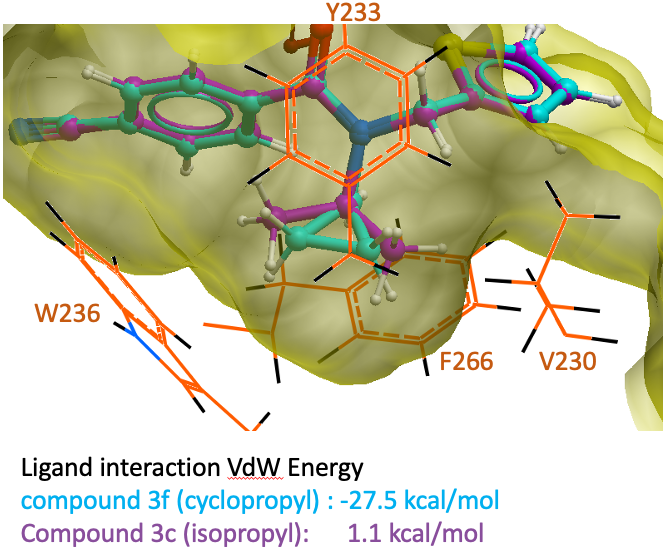

Figure S7. A computational model indicates steric clashes with the isopropyl of compound **3c**. The cyclopropyl of **3f** was replaced with an isopropyl in ICM (Molsoft, San Diego). The isopropyl methyl groups are more distant resulting in increased bulk (the isopropyl geometry is absolutely similar to the one found in ligand bk1, PDB code 3I7B). The energy of the modified flexible ligand was locally minimized in the internal coordinate space, while conserved atoms were tethered to their original position (with a tether weight tzWeight=200). The Van der Waals energy of the bound ligand was calculated in ICM (force-field: protein: ECEPP/3, ligand: mmff).

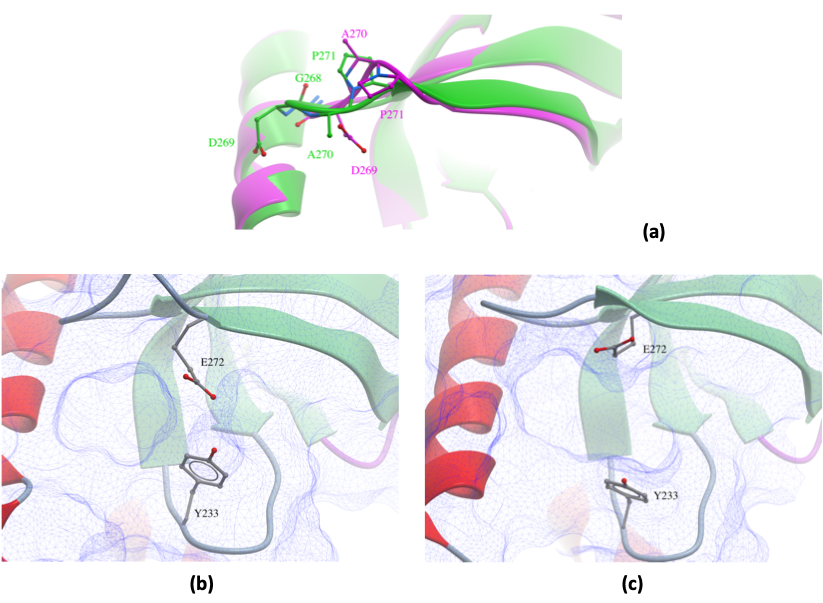

Figure S8. (a) Comparison between the apo (PBD ID: 5VC8, magenta) and bound (green) conformation of the NSD2-PWWP1. The loop residues G268, D269, A270, and P271 connecting the β3 and β4 strands display different conformations in the two structures. (b) In the apo structure the residues E272 and Y233 are closing the pocket. (c) When the ligand binds these residues move away opening the pocket.

**Chemical synthesis of compounds 3c and 3d**

All starting materials were commercially procured and were used without further purification unless specified. Analytical LCMS data for all compounds were acquired using an Agilent 6110 Series system with the UV detector set to 220 nm. Samples were injected (<10 µL) onto an Agilent Eclipse Plus 4.6 × 50 mm, 1.8 µm, C18 column at room temperature. Mobile phases A (H2O + 0.1% acetic acid) and B (MeOH + 0.1% acetic acid) were used with a linear gradient from 10% to 100% B in 5.0 min, followed by a flush at 100% B for another 2 minutes with a flow rate of 1.0 mL/min. Mass spectra (MS) data were acquired in positive ion mode using an Agilent 6110 single quadrupole mass spectrometer with an electrospray ionization (ESI) source. Nuclear Magnetic Resonance (NMR) spectra were recorded on a Varian Mercury spectrometer at 400 MHz for proton (^1^H NMR); chemical shifts are reported in ppm (δ) relative to residual protons in deuterated solvent peaks. Normal phase column chromatography was performed with a Teledyne Isco CombiFlash®Rf using silica RediSep®Rf columns with the UV detector set to 220 nm and 254 nm. The mobile phases used are indicated for each compound. Reverse phase column chromatography was performed with a Teledyne Isco CombiFlash®Rf 200 using C18 RediSep®Rf Gold columns with the UV detector set to 220 nm and 254 nm. Mobile phases of A (H2O + 0.1% TFA) and B (MeCN) were used with default column gradients. Preparative HPLC was performed using an Agilent Prep 1200 series with the UV detector set to 220 nm and 254 nm. Samples were injected onto a Phenomenex Luna 250 x 30 mm, 5 µm, C18 column at room temperature. Mobile phases of A (H2O + 0.1% TFA) and B (MeOH or MeCN) were used with a flow rate of 40 mL/min. A general gradient of 0-15 min increasing from 10 to 100% B, followed by a 100% B flush for another 5 min was used. Small variations in this purification method were made as needed to achieve ideal separation for each compound. Analytical LCMS (at 254 nm) and NMR were used to establish the purity of targeted compounds. All compounds that were evaluated in biochemical and biophysical assays had >95% purity as determined by ^1^H NMR and LC-MS.

**Synthesis of compound 3c**

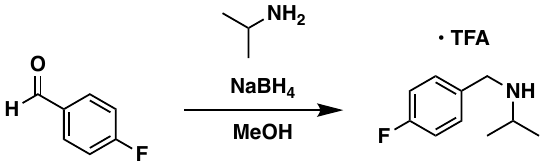

To a flask equipped with a stir bar were added 4-fluorobenzaldehyde (0.25 g, 0.21 mL, 1 Eq, 2.0 mmol) and methanol (10 mL). The flask was sealed with a septum, and isopropylamine (0.59 g, 0.86 mL, 5 Eq, 10 mmol) was introduced *via* syringe. The mixture was stirred at room temperature for 3 hours to allow for imine formation. Next, the flask was cooled in an ice bath, and NaBH_4_ (0.15 g, 2 Eq, 4.0 mmol) was added in one portion. The reaction was allowed to come to room temperature with stirring overnight. The next day, the reaction was diluted with water and extracted 3 times with ethyl acetate. The combined organic layers were washed three times with water and once with brine, then dried over sodium sulfate and concentrated to a clear oil. The oil was taken up in ether (10 mL) and cooled in an ice bath, and TFA (0.34 g, 0.23 mL, 1.5 Eq, 3.0 mmol) was added with vigorous stirring. The voluminous white precipitate formed was filtered off, rinsed with additional ether, and air-dried to provide N-(4-fluorobenzyl)propan-2-amine 2,2,2-trifluoroacetate (111.5 mg, 396.4 µmol, 20 %) as a fluffy white solid.

^1^H NMR (400 MHz, Methanol-*d*_4_) δ 7.54 (dd, *J* = 8.6, 5.4 Hz, 2H), 7.20 (t, *J* = 8.7 Hz, 2H), 4.20 (s, 2H), 3.44 (hept, *J* = 6.5 Hz, 1H), 1.39 (d, *J* = 6.6 Hz, 6H).

LCMS (ESI, +ve mode) expected *m/z* for C10H15FN^+^ [M+H] 168.12, found 168.20

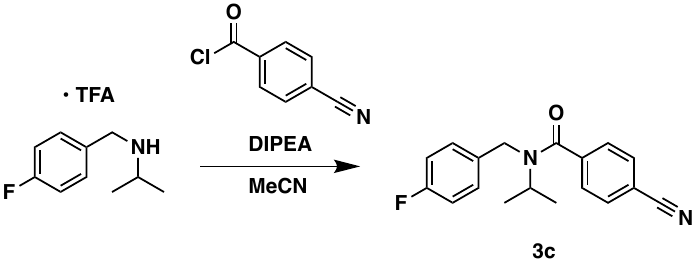

To a vial containing N-(4-fluorobenzyl)propan-2-amine 2,2,2-trifluoroacetate (75 mg, 1.1 Eq, 0.27 mmol) was added acetonitrile (1 mL) and triethylamine (74 mg, 0.10 mL, 3 Eq, 0.73 mmol). The vial was cooled in an ice bath, and 4-cyanobenzoyl chloride (40 mg, 1 Eq, 0.24 mmol) was added in one portion. The reaction was allowed to come to room temperature with stirring overnight. The next day, the reaction was diluted with water and extracted 3 times with ethyl acetate. The combined organic layers were washed once with 0.5 M citric acid, once with water, once with saturated sodium bicarbonate, and once with brine, dried over sodium sulfate, and concentrated to a white solid. Normal phase chromatography over silica gel (0-50% ethyl acetate in hexanes) afforded compound **3c** (47 mg, 0.16 mmol, 65 %) as a clear residue that solidified on standing to a white solid.

^1^H NMR (400 MHz, Methanol-*d*_4_) δ 7.86 (d, *J* = 7.8 Hz, 2H), 7.63 (d, *J* = 7.7 Hz, 2H), 7.41 (s, 2H), 7.07 (t, *J* = 8.6 Hz, 2H), 4.70 (s, 2H), 4.02 – 3.91 (m, 1H), 1.14 (d, *J* = 5.9 Hz, 6H). Major rotamer peaks listed.

LCMS (ESI, +ve mode) expected *m/z* for C18H18FN2O^+^ [M+H] 297.14, found 297.10

**Synthesis of compound 3d**

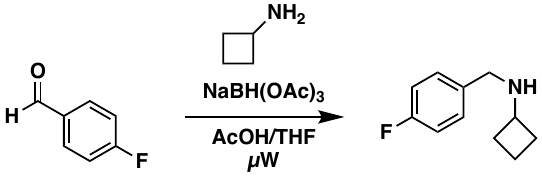

To a flame-dried microwave vial equipped with a stir bar were added 4-fluorobenzaldehyde (100 mg, 86.4 µL, 0.806 mmol) and cyclobutanamine (57.3 mg, 68.8 µL, 0.806 mmol), followed by acetic acid and THF (ratio 1:4; 4ml). The sealed vial was irradiated in the microwave at 60 ˚C (250 W) for 10 min. Upon cooling, Silia*Bond* cyanoborohydride was added (50.6 mg, 0.806 mmol) and stirred at room tempterature for 10 min. The mixture was irradiated in the microwave at 120 ˚C (250 W) for 10 min. The crude mixture was filtered, concentrated under vacuum, and used in the next step without further purification.

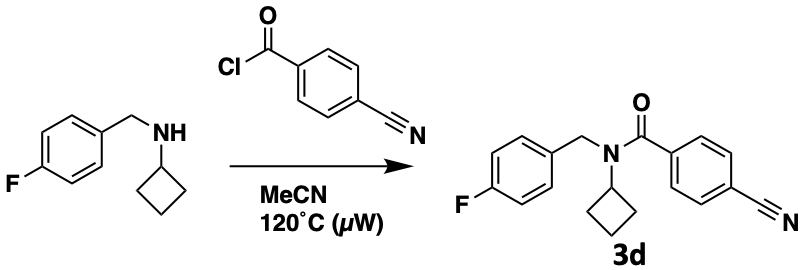

To a flame dried microwave vial equipped with a stir bar were added the N-(4-fluorobenzyl)cyclobutanamine synthesized previously, 4-cyanobenzoyl chloride (100 mg, 0.604 mmol), and acetonitrile (2 mL). The sealed vial was irradiated in the microwave at 120 ˚C for 60 min (250 W). Once the reaction was complete, the volatiles were evaporated. The crude mixture was first purified by normal phase gradient column chromatography (0-100% ethyl acetate in hexanes). The desired fractions were concentrated, re-dissolved in methanol (1 ml), and then purified further by preparative-HPLC ((H_2_O + 0.1% TFA)/MeCN) to afford compound **3d** (56.9 mg, 184 µmol, 30% over 2 steps) as a colorless gum.

^1^H NMR (400 MHz, DMSO-*d*_6_) δ 7.94 (br s, 2H), 7.59 (br s, 2H), 7.33 (br s, 2H), 7.16 (t, *J* = 8.6 Hz, 2H), 4.77 (br s, 2H), 4.06 (br s, 1H), 2.17 – 2.01 (m, 2H), 1.81 (br s, 2H), 1.52 – 1.26 (m, 2H).

LCMS (ESI, +ve mode) expected *m/z* for C19H18FN2O^+^ [M+H] 309.14, found 309.20

**^1^H NMR Spectra**

**4-cyano-N-(4-fluorobenzyl)benzamide**

**Compound 3c**

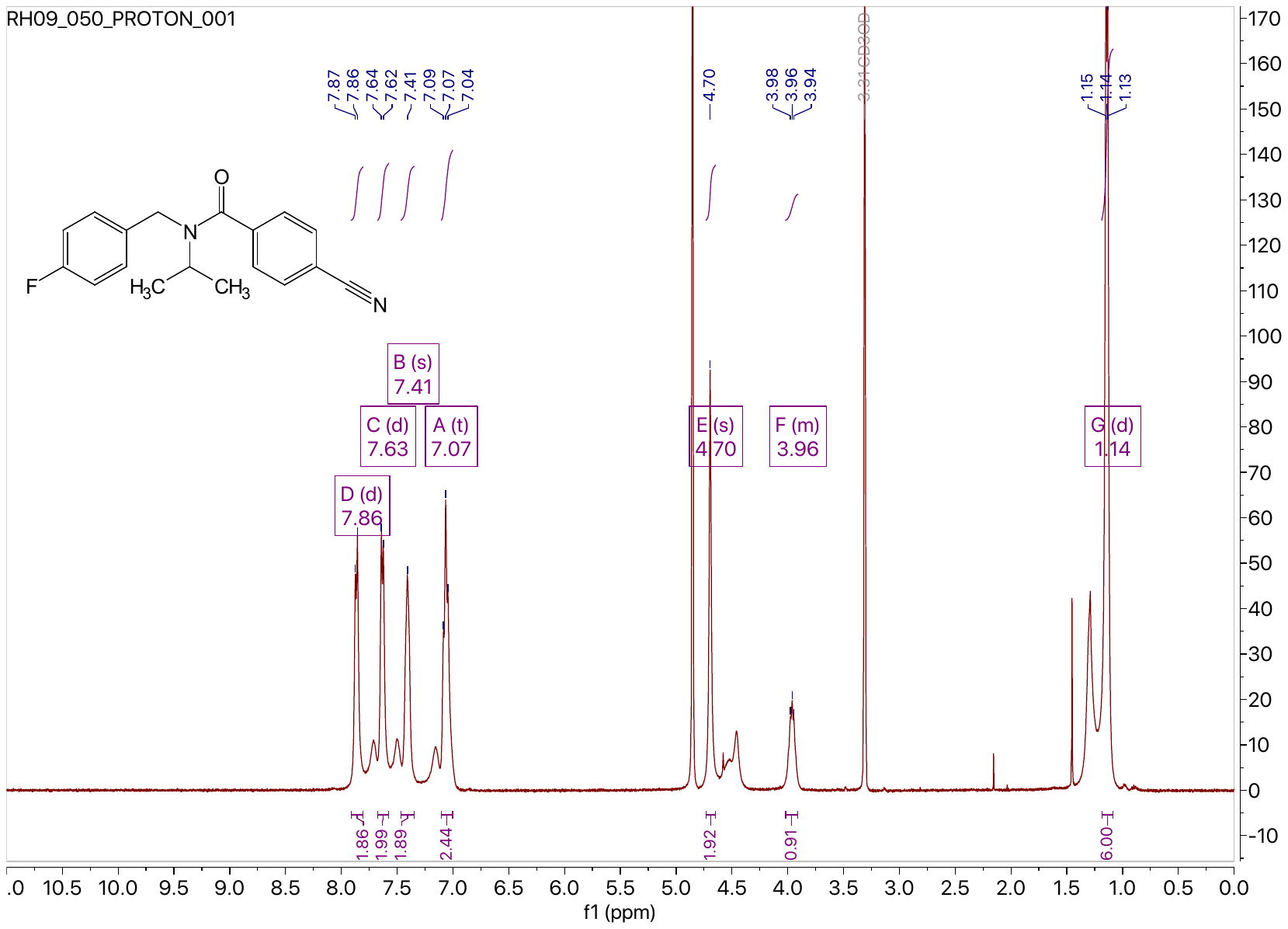

**Compound 3d**

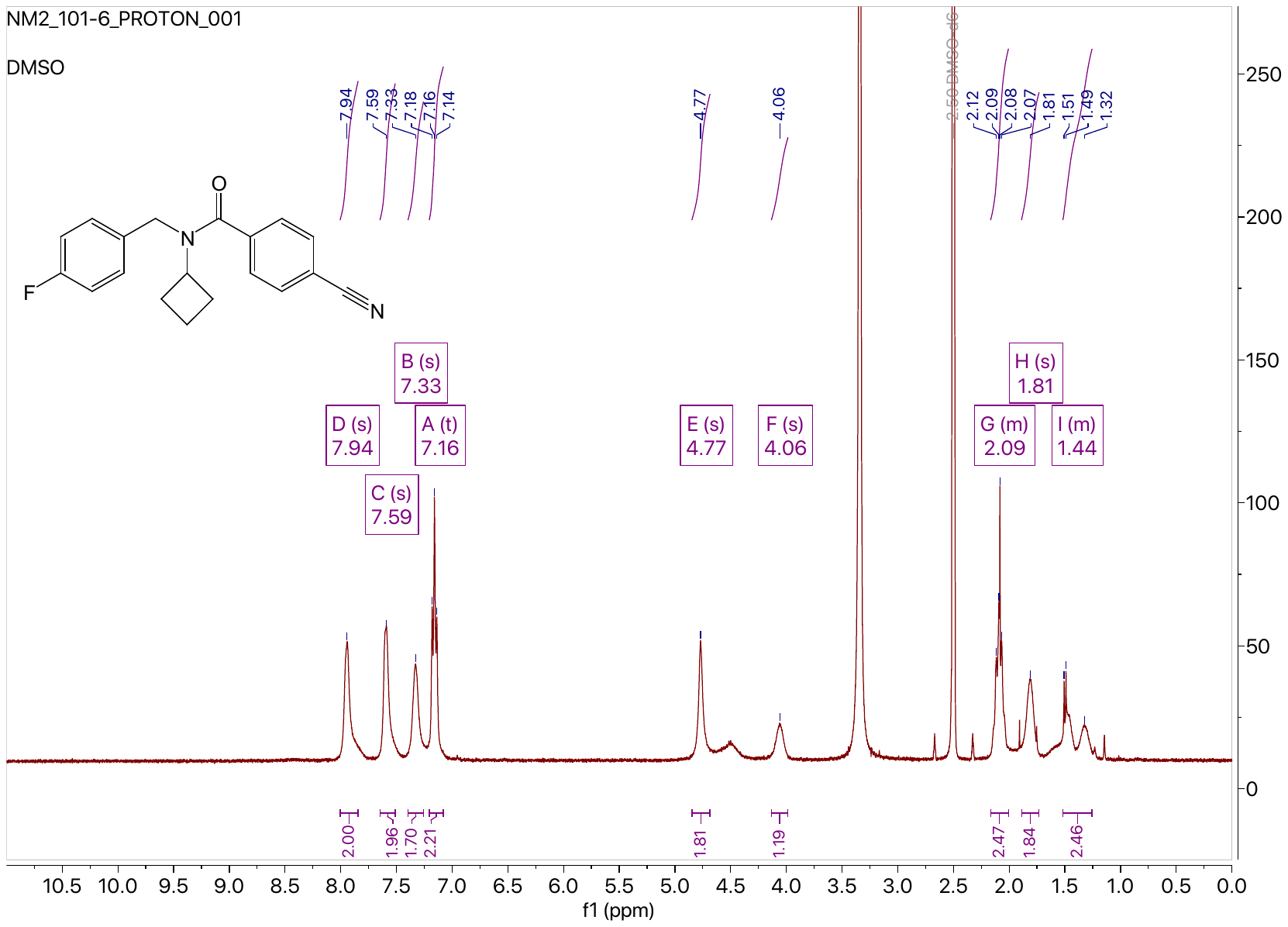
